# Inverse and Postponed Impacts of Extracellular Tau PHF on Astrocytes and Neurons’ Mitochondrial Function

**DOI:** 10.1101/2024.03.19.585791

**Authors:** Valentin Zufferey, Enea Parietti, Aatmika Barve, Jeanne Espourteille, Yvan Varisco, Kerstin Fabbri, Francesca Capotosti, Nicolas Preitner, Kevin Richetin

## Abstract

**Background:** Tauopathies encompass a spectrum of neurodegenerative disorders which are marked by the pathological aggregation of tau protein into paired helical filaments (PHF-tau), neurofibrillary tangles (NFTs) and Glial-fibrillary tangles (GFTs). These aggregates impair cellular, mitochondrial, and synaptic functions. The emergence of extracellular tau (ePHF-tau), featuring a myriad of isoforms and phosphorylation states, presents a challenge in comprehending its nuanced effects on neural cells, particularly concerning synaptic and mitochondrial integrity.

**Methods:** We studied the impact of ePHF-tau (2N4R) on different states and ages of primary cultures of rat neuroglia. Using confocal microscopy and proteomic analysis of synaptosomes, we studied the impact of ePHF-tau on neurite and synapse number. We monitored mitochondrial responses in neurons and astrocytes over 72 hours using advanced fluorescence microscopy for dynamic, high-throughput analysis.

**Results:** Treatment with ePHF-tau has a strong effect on the neurites of immature neurons, but its toxicity is negligible when the neurons are more mature. At the mature stage of their development, we observed a substantial increase in the density of the PSD-95/vGlut1 zone in neurite, suggesting altered synaptic connectivity and ePHF-tau excitotoxicity. Proteomics revealed significant changes in mitochondrial protein in synaptosomes following exposure to ePHF-tau. In the neuronal compartment, real-time imaging revealed rapid and persistent mitochondrial dysfunction, increased ATP production, and reduced mitochondrial turnover. In contrast, we observed increased mitochondrial turnover and filamentation after treatment in the astrocyte processes, indicating cell-specific adaptive responses to ePHF-tau.

**Conclusions:** This study sheds light on the intricate effects of extracellular tau aggregates on neuronal and astrocytic mitochondrial populations, highlighting how tau pathology can lead to mitochondrial disturbances and synaptic alterations. By delineating the differential responses of neurons and astrocytes to ePHF-tau, our findings pave the way for developing targeted therapeutic interventions to mitigate the detrimental impacts of tau aggregates in neurodegenerative diseases.

## Background

Under healthy conditions, the tau protein plays a crucial role in stabilizing microtubules within neurons, which is vital for preserving cellular architecture and facilitating intracellular transport mechanisms [1]. However, as individuals age, the brain may exhibit abnormal accumulations of tau protein, forming insoluble paired helical filaments (PHF-tau). These can further aggregate into neurofibrillary tangles (NFTs), which are composed of various phosphorylated tau species. This phenomenon underpins the pathology of a diverse group of disorders known as tauopathies, which encompass more than 20 diseases, such as Alzheimer’s disease (A.D.), progressive supranuclear palsy (PSP), and frontotemporal dementia (FTD) [2]. Despite the common feature of tau accumulation in tauopathies, significant distinctions exist regarding the distribution of tau pathology, the occurrence of tau inclusions in glial cells, the association with other neuropathological features such as amyloid deposits or Lewy bodies, and the specific tau isoforms and aggregates present [3].

Much research has focused on the deleterious effects of intracellular Tau aggregation, particularly concerning synaptic functions and mitochondrial integrity. Various tau species, including extracellular NFTs and PHF-tau (ePHF-tau), can be released into the extracellular space during neurodegenerative states. This process of cell-to-cell transmission has been thoroughly investigated, revealing how it facilitates the spread of tau pathology by compromising the function of endogenous tau in neighboring neurons [4,5]. Studies have shown that extracellular paired helical filaments (PHFs) can trigger extensive tau accumulation within neurites, mainly affecting postmitotic neurons [6]. It has also been suggested that PHFs indirectly interfere with dynein/kinesin-mediated axonal transport, underscoring the intricate impact of tau on neuronal function [7]. The wide variability of extracellular tau, including differences in isoform and phosphorylation levels, complicates our understanding of its exact pathophysiological consequences on synaptic structures.

Although most of the related work has focused on understanding the deleterious effects of various forms of tau on neurons, several studies have recently demonstrated that glial cells, particularly astrocytes, are capable of accumulating tau in its hyperphosphorylated and aggregated form in different contexts and pathologies. In this context, we previously demonstrated that the mitochondrial system is particularly susceptible to the different forms of 3R and 4R Tau that accumulate or recover [8,9]. To further extend our understanding of the cellular and mitochondrial impacts of Tau pathology, we explored the impact of ePHF aggregates on 2D pure neuron rat primary cultures and 3D neuro-glia rat primary cultures. We aimed to decipher the mechanisms by which ePHF influences cell function through an integrative approach combining confocal imaging, real-time fluorescence imaging and proteomics analysis of synaptosomes. We shed light on the complexity and temporality of the effect of ePHFs on different synaptic compartments, with a particular focus on the mitochondrial system of neurites and astrocytic processes. This work reveals the subtle perturbations induced by ePHF-tau and its ability to modulate synaptic architecture through its differential effects on the mitochondria of neurons and astrocytes.

## Methods

### Isolation of human brain derived PHF

Tau paired-helical filaments (PHF) were isolated from post-mortem frontal cortex tissue of an Alzheimer’s disease (A.D.) patient (purchased from Tissue Solution). The enrichment procedure was modified from Jicha et al, 1997 and Rostagno and Ghiso, 2009. Briefly, approximately 15 g of frontal cortex tissue were thawed on ice and homogenized using a glass Dounce homogenizer in 50 mL of a homogenization buffer consisting of 0.75 M NaCl, 100 mM (MES) pH 6.8, 1 mM EGTA, 0.5 mM MgSO4, 2 mM DTT, and protease inhibitors (Complete; Roche 11697498001). The homogenate was then incubated at 4°C for 20 min for microtubule depolymerization, before being transferred into polycarbonate centrifuge bottles (16 x 76 mm; Beckman 355603) and centrifuged at 11,000 g (12,700 RPM) in an ultracentrifuge (Beckman, XL100K) for 20 min at 4°C using the pre-cooled 70.1 rotor (Beckman, 342184). Pellets were kept on ice. Supernatants were pooled into polycarbonate bottles and centrifuged again at 100,000 g (38,000 RPM) for 1 h at 4°C in the 70.1 Ti rotor to isolate PHF-rich pellets, whereas soluble tau remained in the supernatants. The pellets from the first and second centrifugations were resuspended in 120 mL of extraction buffer [10 mM Tris-HCl pH 7.4, 10% sucrose, 0.85 M NaCl, 1% protease inhibitor (Calbiochem 539131), 1 mM EGTA, 1% phosphatase inhibitor (Sigma P5726 and P0044)]. The solution was then transferred into polycarbonate centrifuge bottles (16 x 76 mm; Beckman 355603) and centrifuged at 15,000 g (14,800 RPM) in an ultracentrifuge (Beckman, XL100K) for 20 min at 4°C using the 70.1 Ti rotor. In the presence of 10% sucrose and at low-speed centrifugation, most PHF remained in the supernatant whereas intact or fragmented NFTs and larger PHF aggregates were pelleted. The pellets were discarded. 30% Sarkosyl (Sigma L7414-10ML) was added to the supernatants to a final concentration of 1% and stirred at R.T. for 1 h. This solution was then centrifuged in polycarbonate bottles at 100,000 g (38,000 RPM) for 1 h at 4°C in the 70.1 Ti rotor, and the pellets containing PHF-rich material were resuspended in 50ul PBS/ 1g of brain tissue. The resuspended PHF was then sonicated on ice for 60s with 1s-on/1s-off cycles at 20% amplitude. Aliquots were snap frozen and stored at −80°C.

### Preparation of Tau aggregates

Tau PHF isolated from Human A.D. brain was used as seed to induce the templated aggregation of human full length 2N4R Tau monomers (35uM, Biotechne in a 72h incubation at 37C on a shaker at 1000 rpm. These aggregates were stored at −20°C. Three days before the treatment of neurons, these aggregates were thawed and used as seeds (1:200) in a secondary aggregation reaction in the presence of 10 µM tau monomers (72h at 37C, 1000 rpm).

### Lentiviral vector production and infection

The L.V.s were pseudotyped as vesicular stomatitis virus glycoprotein G (VSV-G). LV-PGK-MitoTimer and LV-PGK-MitoGoTeam2 (G1B3 promoter instead of PGK for astrocytes) were produced by transfection of human embryonic kidney (HEK) *293T* cells (mycoplasma-negative, ATCC, LGC Standards GmbH, Germany). L.V.s were concentrated by ultracentrifugation and resuspended in phosphate-buffered saline (dPBS, Gibco, Life Technologies, Zug, Switzerland) supplemented with 1% bovine serum albumin (BSA, Sigma_Aldrich, Buchs, Switzerland). The viral particle content in each batch was determined using a p24 antigen enzyme-linked immunosorbent assay (p24 ELISA, RETROtek; Kampenhout, Belgium). The stocks were stored at −80°C until use and diluted to 100,000 ng/ml in PBS/1% BSA. For neuronal expression of mitochondrial biosensors, cells were infected at DIV 5 with lentiviral vectors (L.V.s) expressing reporter genes, mainly in neurons. The LV-PGK vectors used in this study have been previously described by Schwab and colleagues [10]. The reporter gene contained a woodchuck hepatitis virus B postregulatory element (WPRE) and the mouse PGK promoter. Astrocytes were infected at 8 DIV with lentiviral vectors (LV) expressing the mitochondrial reporter gene. The gfaABC1D promoter was ligated to enhancer B(3) to generate the G1B3 promoter, which was subsequently cloned and inserted into the SIN-cPPT-gateway-WPRE-miR124T transfer plasmid, which contains four copies of the neuron-specific miRNA-124 target sequence (miR124T; full homology) to repress transgene expression in neurons, WPRE and the central polypurin tract (cPPT) to increase transgene expression and a 400-nucleotide deletion in the long 3′ terminal repeat (self-inactivating vector) to increase biosafety.

### 2D Primary pure neuron culture

Rat primary cortical cultures were prepared from Sprague-Dawley rats (Charles River Laboratories L’Arbresle, France) at postnatal day 1. The neocortex and hippocampus were isolated by removing the dura, brainstem, olfactory bulbs and subcortical regions. Isolated cortex and hippocampus were cut into small pieces and dissociated by enzymatic digestion for 30 minutes at 37°C in dissociation buffer (papaine, CaCl2, EDTA, and HEPES; all from Invitrogen, Carlsbad, CA, USA). DNase (Invitrogen, Basel, Switzerland) was added, and the mix was incubated for another 30 min at 37C. After mechanical dissociation consisting in pipetting the tissue up and down, the cell suspension was passed through a cell strainer and dispersed cells were plated onto 96-well tissue culture plates coated with poly-D-Lysine (Corning, 3841) at 30K cells/well. Cells were plated in Neurobasal medium (Invitrogen, Basel, Switzerland) without phenol red, with the addition of L-glutamine (2 mM; Sigma, Buchs, Switzerland), 10% FCS (Invitrogen, Basel, Switzerland), and penicillin/streptomycin (Sigma, Buchs, Switzerland) and incubated at 37C in the presence of 5% CO2. 1.5 hours after plating the cells, the medium was replaced with Neurobasal medium containing 2mM L-Glutamine, B-27TM (Gibco, 17504-044), and penicillin/streptomycin. After 4 days in culture, cell proliferation was blocked by treatment with 2.5 µM cytosine arabinoside (Invitrogen, Basel, Switzerland).

### 3D Primary neuron-glial cultures

For 3D culture, hippocampus of E17 Wistar rat embryos was dissected, and the cells were dissociated with a neuronal dissociation kit (130-092-628). Mixed cells were plated at a density of 30K cells/cm2 in 24-well or 96-well glass-bottom plates (Ibidi 82426, Corning™ 356234) coated with Matrigel in 200 µM DMEM (Gibco 41965-039, 25 mM glucose) supplemented with 0.25% L-glutamine (Gibco, 25030081), 1% penicillin/streptomycin (Thermo Fisher Scientific 15140-122) and 2% B27 (17504044, Gibco) and incubated at 37°C and 5% CO2.

### Immunofluorescent stainings

Three days after treatment, cells were fixed with 4% PFA (Sigma, 158127-500G) for 15 min at 37°C and stored in PBS at 4°C for further analysis. All the incubations and extractions were carried out in PBST (10010-015, Gibco), 0.3% Triton X-100 (Sigma–Aldrich, X100-100ML), and 3% NHS (Gibco, 16050-122). Primary antibody incubation was performed overnight with rabbit anti-VGlut1 (Cell Signaling Technology, 1230.31; 1:500), chicken anti MAP2 (Abcam, ab5392, 1:1’000), goat anti-PSD95 (Abcam, ab12093; 1:500), or mouse anti-NeuN (Chemicon, MAB337) and anti-GFAP (Dako, GA52461-2) antibodies. Secondary antibody incubation was conducted for 60_min at room temperature with Alexa Fluor 488-, 555- or 647-conjugated highly cross-adsorbed donkey anti-goat, donkey anti-rabbit or donkey anti-chicken antibodies, respectively (1:500; Invitrogen A31573, Invitrogen A11055, Jackson IR 703-545-155). After additional incubation in DAPI (Merck, 268298; 1:5,000), the cells were stored at 4_°C in PBS.

### Microscopy and image acquisition

In this study, 2D cultures (Fig.1) and live imaging (Fig. 4-5) were acquired with an inverted microscope (Nikon Eclipse Ti-2) equipped with an Okolab cage incubator (H201-T-UNIT), the Perfect Focus System (PFS), CFI Plan Apochromat Lambda D 40X and 60X Oil, and Lumencor SPECTRA X Light Engine. 3D culture (Fig.2) were acquired with fully motorized model Ni-E equipped with spinning-disk confocal technology (CrestOptic, X-Light V3) and CFI Plan Apochromat Lambda D 40X and 60X Oil. All images were acquired with Nikon’s software suite NIS HC v.5.42.

**Figure 1:**
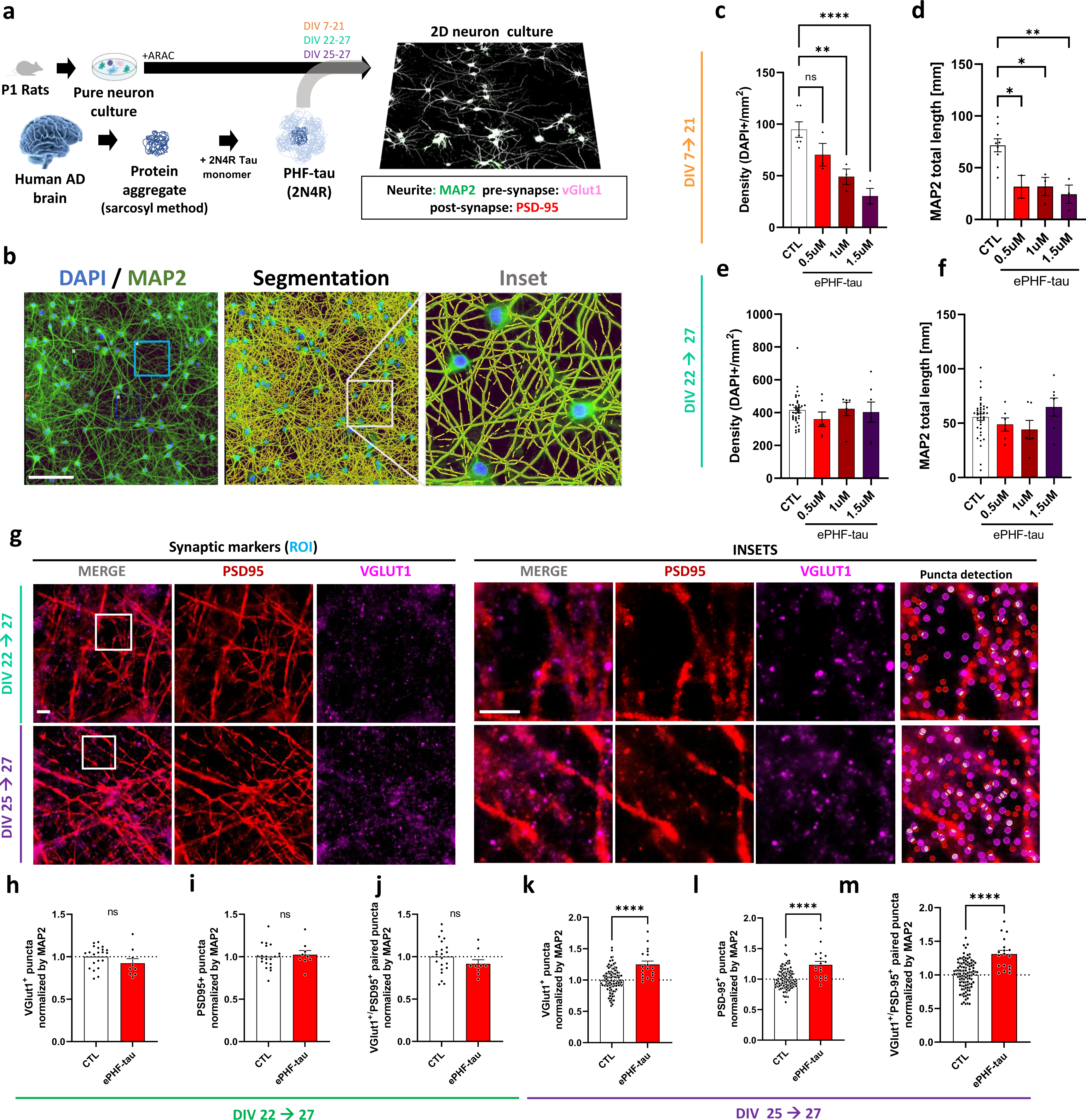
ePHF-tau affects neurites development and synapse maturation. (a) Scheme of the experimental design for tau aggregate treatment in pure neuron cultures. (b) Illustration of a fluorescence micrographs of dapi (DNA) and map2 (neurites) staining, and of the segmentation of neurites and nuclei. (blue squares indicates an examples ROI for subsequent synapses analysis). (c-f) Bar plots of the quantifications of cell numbers (c, e) and neurites total length (d, f), respectively with early or late treatment. (g) Examples ROI image used for synapses marker analysis (PSD-95, vGlut1). White squares correspond to the insets shown on the right, in which examples of the segmentation used for analysis are depicted. (h-J) Bar plots of the control normalized counts of pre-synaptic (h), post-synaptic (i) and associated markers (j) (medians) with early treatment. (k-m) Same as (h-j) for later treatment. (N_=_3 cultures, 2 replicates per conditions, 1 large image per replicate, and 3 ROIs per image for synapse analysis; **** p <0.0001).

**Figure 2:**
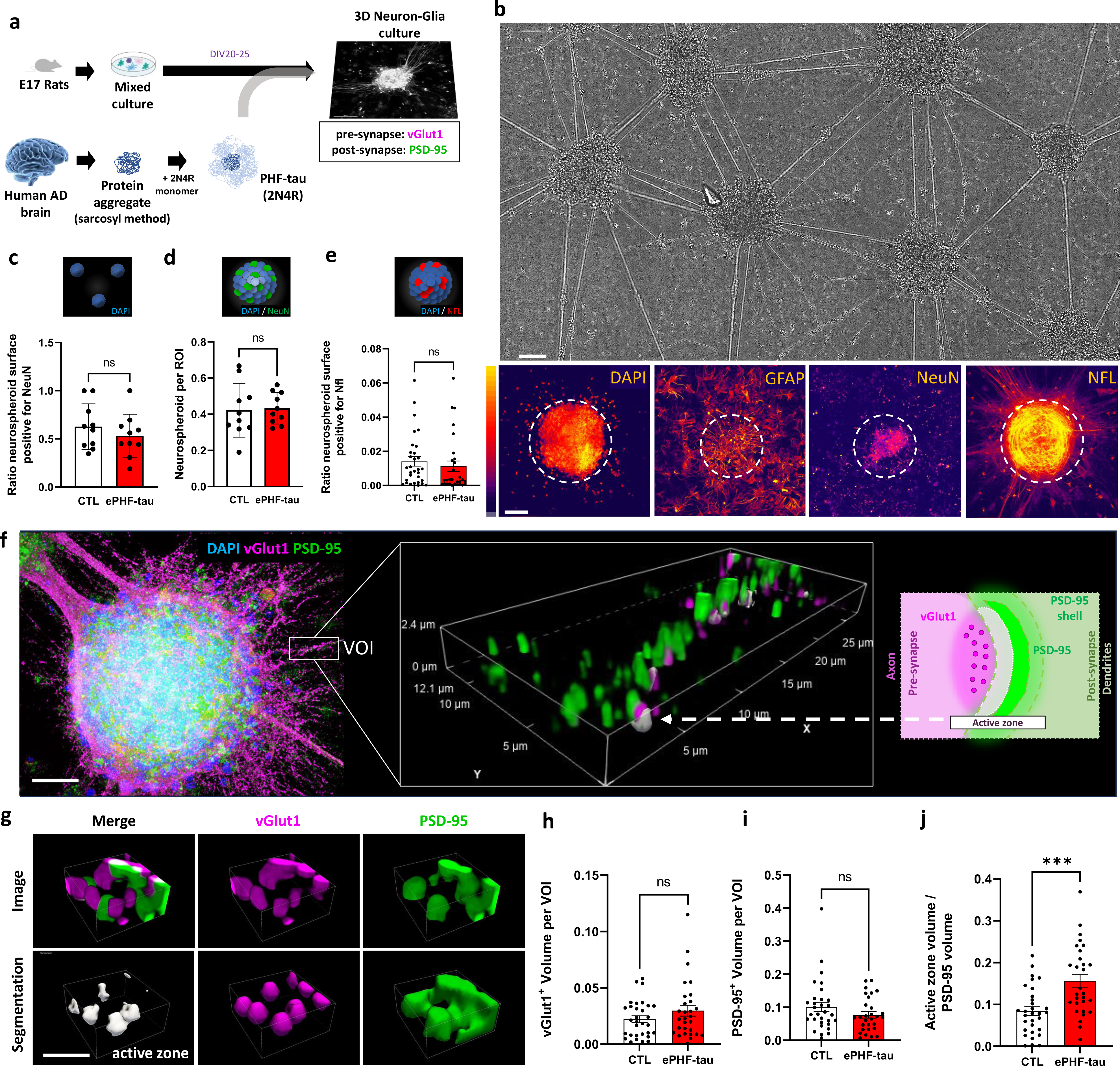
ePHF-tau induces modifications in synaptic markers in mixed cultures. (a) Scheme of the experimental design for tau aggregate treatment in mixed cultures. (b) Micrographs of a large section of culture depicting neurospheroids in bighfield. Examples of stainings for nucleis, glial and neuronal markers (Dapi, GFAP, NeuN, NFL). (c-e) Bar plots of the quantification of neuropheroid density (c), as well as NeuN (d) and neurofilament (e) coverage of neurospheroids. (f) Illustration of neurospheroid stained for synaptic markers with example of an analyzed volume depicted with a white rectangle and illustrated in the center. Overlayed white objects (e.g. white arrow) correspond to the identified active zones. On the right, a small diagram illustrating how the active zone object were reconstructed from PSD-95 and vGlut1 signals. (g) Detailed examples of pre and post synaptic markers images (top) and associated 3D reconstruction (bottom). (h-j) Bar plots of the total PSD-95 (h) and vGlut1 (i) volumes (medians), and active zone volume per surface of PSD95. (N_=_3 cultures, total of 25 neurospheroids scanned, and 5 neurites VOIs analyzed per image; *** p <0.001).

**Figure 3:**
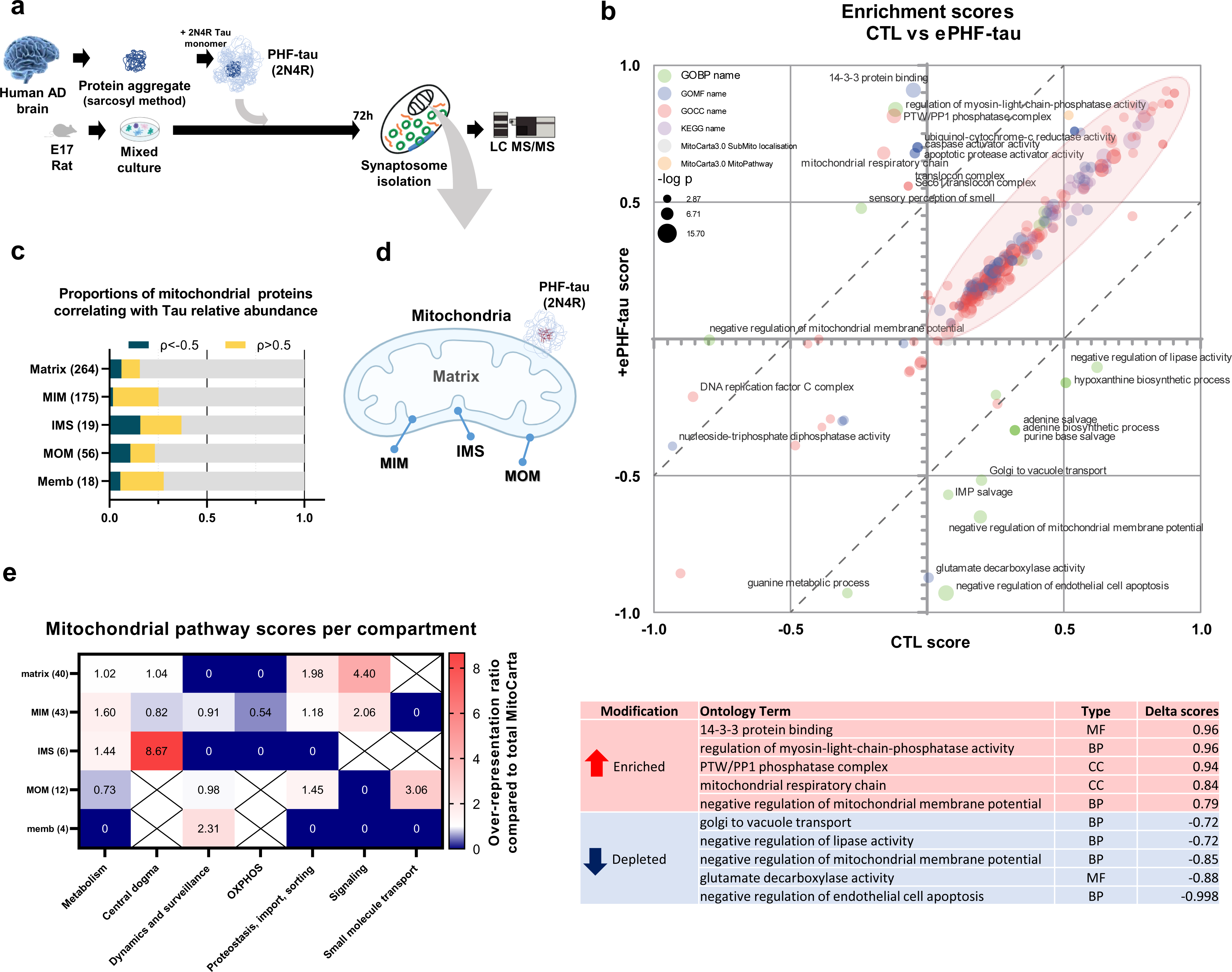
Effect of ePHF-tau on synaptic mitochondrial composition. (a) Scheme of the experimental design for synaptosome proteomics. (b) Two-dimensional (2D) annotation term enrichment analysis using LFQ as a quantitative indicator of protein abundance. Enrichment scores in the CTRL and ePHF-tau groups are shown on the X- and Y-axes, respectively. The table shows the enriched terms with the top five score differences (red: enriched with tau, blue: depleted). (c) Stacked bar plot of the distribution of mitochondrial proteins in each sublocalization, grouped by the values of their correlation with tau protein levels. (d) Scheme of a mitochondrion labeled with the Mitocarta 3.0 annotations of sublocalization. (e) Heatmap of the over-/underrepresentation of pathways in each subcompartment for mitochondrial-correlated proteins when normalized to the proportions obtained from the total MitoCarta database. Crossed-out cells indicated that these ontologies did not occur in Mitocarta. (N_=_3 cultures per condition).

**Figure 4:**
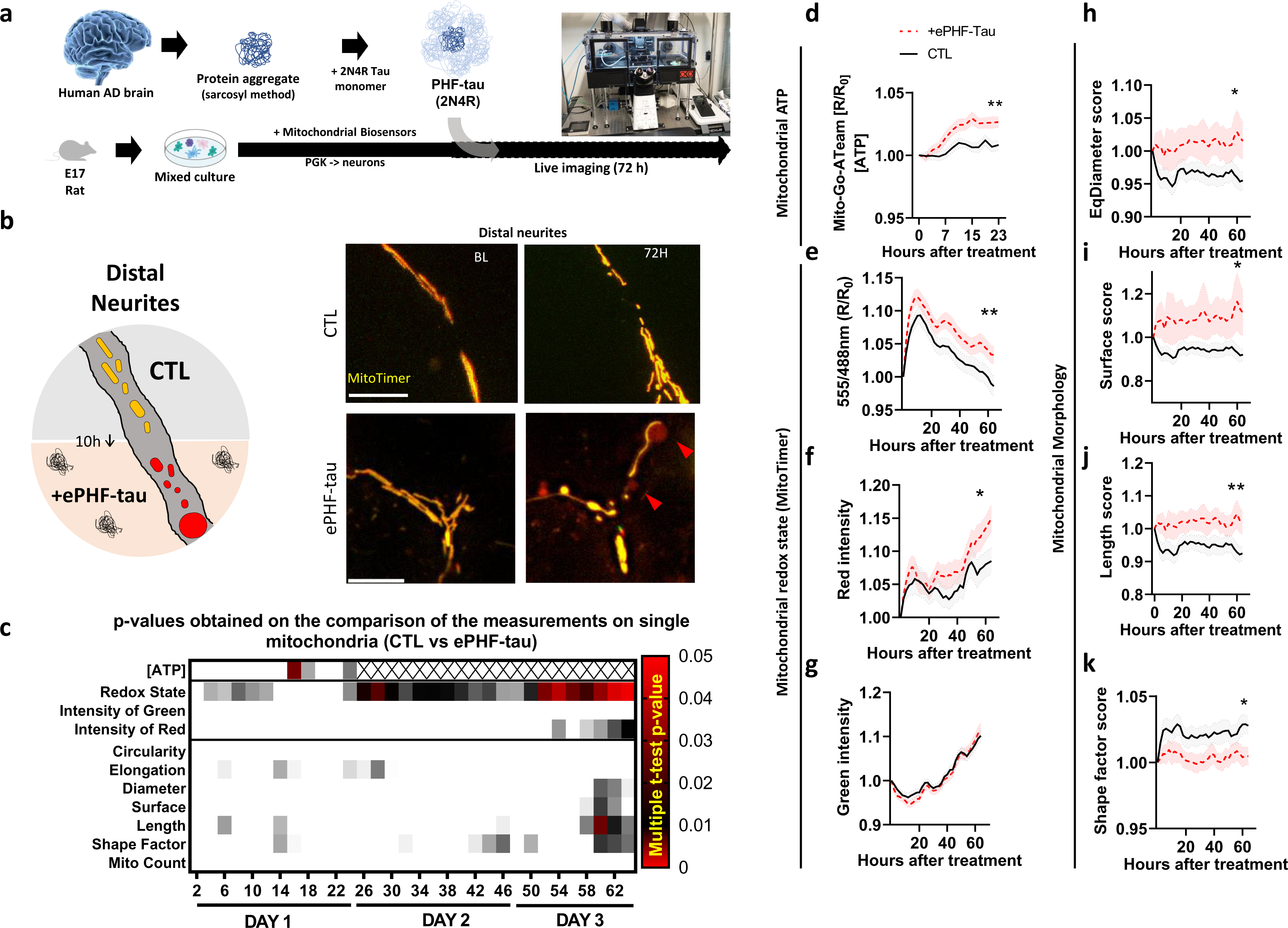
ePHF-tau negatively affects mitochondrial turnover in neurites. (a) Scheme of the experimental design: primary mixed cultures were equipped with mitochondrial sensors and imaged over time in the presence of ePHF-tau. (b) Fluorescence microscopy images of Mitotimer in neurites of the CTRL and ePHF-tau treatment groups at baseline, 24 h and 72 h and a diagram of the analyzed region and the observed mitochondrial effect. (c) Heatmap of p values obtained for the comparison of scores between the CTRL and ePHF-tau treatment groups over time (0-72 h) (multiple t tests; red values indicate p values<0.05). (d-k) Line plots of time series showing significant differences in the mitochondrial ATP content concerning the Mito-GoATeam2 ratio (R/R0, up to 24 h). (d) Mitotimer red (555 nm)/green (488 nm) redox state ratio (e) red (555 nm) fluorescence intensity (f) and green (488 nm) fluorescence intensity (g) and mitochondrial morphology with diameter (h), surface (i), length (j), and shape factor (k). (N_= 3_cultures; 24 neurites analyzed per culture, *: p value<0.05, **: p value<0.01.)

**Figure 5:**
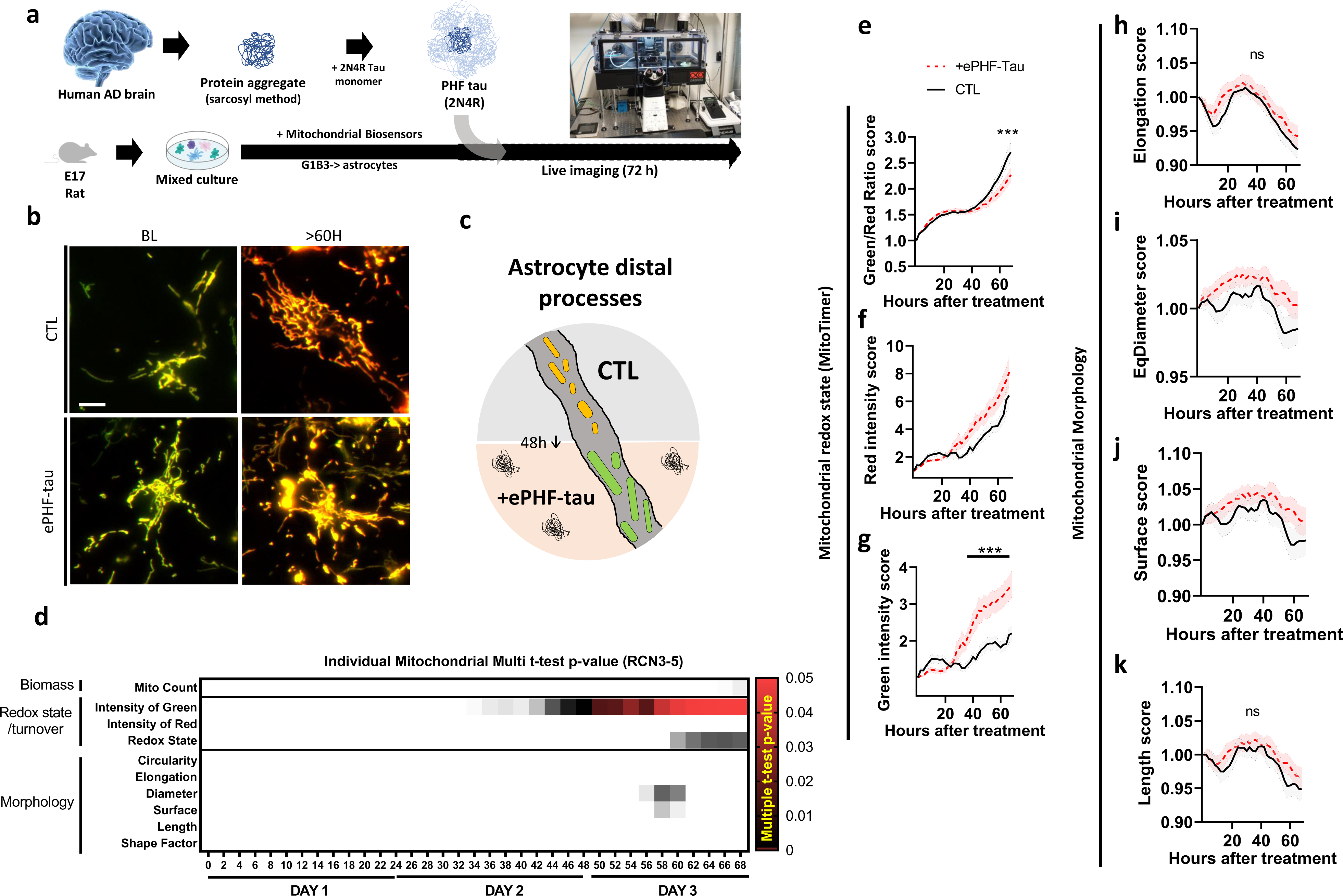
ePHF-tau positively affects mitochondrial turnover in astrocytes. (a) Scheme of the experimental design: primary mixed cultures were equipped with specific mitochondrial sensors and imaged over time in the presence of ePHF-tau. (b) Fluorescence microscopy images of astrocyte mitochondria from the CTRL and ePHF-tau treatment groups at baseline and late time points. Single mitochondria on cellular extensions were sampled for analysis. The diagram summarizes the observations. (d) Heatmap of p values obtained for the comparison of scores between the CTRL and ePHF-tau treatment groups over time (0-72 h) (multiple t tests; red values indicate p values<0.05). (e-k) Line plots of time series showing significant differences in mitochondrial features mesures with MitoTimer in astrocytes. (e) Mitotimer Red (555 nm)/Green (488 nm) redox state ratio. (f) Red (555 nm) fluorescence intensity. (g) Green (488 nm) fluorescence intensity of MitoTime. Mitochondrial morphology with diameter (h), surface (i), length (j), and shape factor (k). (N_=_3 cultures; 140-150 cells analyzed per culture), *: p value<0.05, **: p value<0.01.

### Images analysis

Images were analyzed using Nikon NIS AI or H.C. v.5.42 and the General analysis 3 (GA3) module of this software. For 2D culture analysis, 8000×8000 pixels mosaic images of each well were analyzed to quantify the number of DAPI-labeled cells and neuritic segments, using MAP2 labeling reconstruction. MAP2 was segmented and skeletonized to measure total branch length. MAP2 in the vicinity of DAPI+ nuclei was excluded by subtraction of dilated the DAPI objects.

For synapse analysis all images were enhanced AI-denoising tool. VGlut1+ and PSD-95+ labeled puncta were quantified in regions of interest free of cell nuclei. Synaptic markers were analyzed using a spot detection algorithm. Overlay points between spots were quantified to detect vGlut1/PSD-95 active areas. All data were normalized by the MAP2 area per ROI. The GA3 analysis script used is accessible in the supplementary material. For the analysis of 3D cultures, confocal acquisitions at 60X were performed on approximately 30 focal sections of 0.3 µm of 2048×2048 pixels. Between 5 and 10 volumes of interest located in proximal neurites emanating from neurospheroids were selected to quantify vGlut1 and PSD95 in three dimensions. Volumes corresponding to PSD95 were used to create three-dimensional objects, while a spot detection algorithm was used to identify vGlut1 puncta. To include only vGlut1 volumes in contact with PSD-95, the PSD95 volume was dilated by 0.1 µm. The sum of the volumes of vGlut1 spots in the dilated volume of PSD95 (PSD-95 shell) was then calculated. All GA3 analysis scripts are available in the data repository.

Mitochondrial live monitoring of morphology and turnover was conducted as previously described [8,9] at 40x magnification. We acquired images before treatment as a baseline (B.L) and then every 2 h for 3 days. Live Mitotimer analysis was conducted within a small ROI placed on distal cell processes to ensure the selection of cellular endpoints. A minimum of 12 processes were analyzed for each image sequence. Briefly, Total mitotimer was obtained from the two fluorescent channels and used for segmentation. Only individual and well reconstructed mitochondria were considered for measurements of morphology and fluorescence intensity values (Red and Green intensities, red/green ratio, elongation, area, length, skeleton branches, junctions, circularity, diameter, surface, shape factor, and mitochondria number). Finally, after log transformation and averaging by frame, a score was calculated a the fold-change relatively to the first frame (baseline). To analyze mitochondrial ATP levels in neurons with neuronal Go-ATeam2 [11] expression, single mitochondrial units were semi manually segmented into 12 axons per replicate to conduct ratio measurements. The measurements were further processed like Mitotimer.

### Synaptosome extraction

The cells were rinsed several times, collected, and snap-frozen. Homogenization was performed on ice in PBS. The samples were centrifuged at 1300 × g for 3 min at 4° C to pull down the membrane fragments, nuclei, and cells. The supernatant was coupled with Anti-tomm22 micro-Beats (30min at 4° C) to remove free mitochondria through Magnetic-assisted cell sorting (MACS) (Miltenyi Biotec, 130-096-946, 130-042-401). The mitochondria-depleted flow-through was centrifuged at 13,000 × g for 10 min at 4° C to pellet synaptosomes (100ul of PBS resuspension, stored frozen until further analysis).

### Synaptosomes proteomic mass spectrometry

**(1)** Protein digestion: Synaptosomal fractions were digested according to a modified version of the iST method 80 (named miST method). Briefly, 50 μl solution of PBS was supplemented with 50 μl miST lysis buffer (1% sodium deoxycholate, 100 mM Tris pH 8.6, 10 mM DTT) and heated at 95°C for 5 min. Samples were then diluted 1:1 (v:v) with water, and reduced disulfides were alkylated by adding ¼ vol of 160 mM chloroacetamide (final 32 mM) and incubating at 25°C for 45 min in the dark. Samples were adjusted to 3 mM EDTA and digested with 0.5 μg Trypsin/LysC mix (Promega #V5073) for 1h at 37°C, followed by a second 1h digestion with a second and identical aliquot of proteases. To remove sodium deoxycholate and desalt peptides, two sample volumes of isopropanol containing 1% TFA were added to the digests, and the samples were desalted on a strong cation exchange (SCX) plate (Oasis MCX; Waters Corp., Milford, MA, USA) by centrifugation. After washing with isopropanol/1%TFA, peptides were eluted in 250 μl of 80% MeCN, 19% water, and 1% (v/v) ammonia; **(2)** Liquid chromatography-tandem mass spectrometry: Eluates after SCX desalting were frozen, dried, and resuspended in variable volumes of 0.05% trifluoroacetic acid and 2% acetonitrile to equilibrate concentrations. Approximately 1 μg of each sample was injected into the column for nanoLC-MS analysis. **(3)** M.S. and M.S. data analysis: Dependent LC-MS/MS analysis of the TMT sample was performed on a Fusion Tribrid Orbitrap mass spectrometer (Thermo Fisher Scientific) interfaced through a nano-electrospray ion source to an Ultimate 3000 RSLCnano HPLC system (Dionex). Peptides were separated on a reversed-phase custom-packed 40 cm C18 column (75 μm ID, 100 Å, Reprosil Pur 1.9 μm particles, Dr. Maisch, Germany) with a 4-76% acetonitrile gradient in 0.1% formic acid (total time 140 min). Full MS survey scans were performed at 120’000 resolution. A data-dependent acquisition method controlled by the Xcalibur 4.2 software (Thermo Fisher Scientific) was used to optimize the number of precursors selected (“top speed”) of charge 2+ to 5+ while maintaining a fixed scan cycle of 1.5s. The precursor isolation window used was 0.7 Th. Full survey scans were performed at a 120’000 resolution, and a top speed precursor selection strategy was applied to maximize acquisition of peptide tandem M.S. spectra with a maximum cycle time of 0.6s. HCD fragmentation mode was used at a normalized collision energy of 32% with a precursor isolation window of 1.6 m/z, andthe MS/MS spectra were acquired in the ion trap. The peptides selected for MS/MS were excluded from further fragmentation for 60s. Tandem MS data were processed using MaxQuant software (version 1.6.3.4) 81 incorporating the Andromeda search engine 82. UniProt reference proteome (RefProt) databases for Homo sapiens and mice were used, supplemented with sequences of common contaminants. Trypsin (cleavage at K, R) was used as the enzyme definition, allowing for two missed cleavages. The carbamidomethylation of cysteine was specified as a fixed modification. N-terminal acetylation of proteins and oxidation of methionine are specified as variable modifications.

### Proteomic data analysis

MaxQuant data were further processed using the Perseus software[12] and Microsoft Excel. We used a cutoff for the presence of a protein in a sample such as Razor Peptide Score >2 and MS/MS Count >2; 29% of the proteins were discarded in this process. IBAQ values are the sum of the intensities of all unique peptides for a protein divided by the number of theoretical tryptic peptides between six and 30 amino acids in length[17]. LFQ were calculated according to the number of unique peptides of a protein on the total number of peptides. The relative protein abundance was calculated as the sum of the relative intensity-based absolute quantification (rIBAQ) of all selected proteins that were independently present in each replicate. Protein Annotation was conducted in Perseus using the Gene Ontology databases (Biological Process (B.P.), Cellular Compartment (CC), Molecular Function (M.F.)) [13,14] and MitoCarta3.0 [15,16] databases for mitochondrial protein localization and pathway ontologies. 2D enrichment analysis of annotations was performed in Perseus to assess synaptosomal extraction quality and identify differentially enriched ontologies. We used the MitoCarta annotation to filter the mitochondrial proteins and generate subsets per localization. We conducted a Pearson correlation analysis of the LFQs of mitochondrial proteins in relation to tau protein. We then selected the results with a cutoff of 0.5 on the absolute Pearson coefficient to form groups of positively and negatively correlating (or uncorrelated) proteins. The overrepresentation of pathways at each mitochondrial localization was estimated as the fold-change between the pathway distribution observed from the total Mitocarta database and the distribution observed in the mitochondrial protein subset.

### Statistical analysis

Values are presented as the mean ± SEM; Statistical analyses were performed on raw data with Graphpad Prism software, Shapiro-Wilk tests were performed to test distribution normality. The level of significance was set to P < 0.05. Mitochondrial biosensor data were log-transformed, and multiple t-tests for control versus ePHF conditions were performed at each time point. Mass spectrometry data analysis: 2D annotation enrichment analysis was performed using Benjamini-Hochberg FDR Truncation with a 2% cutoff[18]. Pearson’s correlation analysis was used to evaluate the correlation of proteins MAPT amounts. A cutoff with a correlation coefficient ≥0.5 was used to select correlating proteins for further analysis. Synapse active zone analysis: Comparison of counts, surfaces, and volumes was performed with t-tests or Mann-Whitney, respectively, for normal and non-normal distribution of data. Immunohistochemical analysis: one-way ANOVA of optical densities and mitochondrial data.

## Results

### Impact of ePHF-tau on neurites and synapse development

In this study, we first isolated PHF-tau from the insoluble sarkosylated fraction extracted from the frontal cortex of Alzheimer’s patients. This human PHF-tau was combined with a high amount of monomeric 2N4R tau to generate PHF-tau(2N4R), subsequently referred to as ePHF-tau, following the methodology developed by Courade et al [19]. To further our understanding of the impact of ePHF on neurites and synapses, pure neuronal cultures were exposed to various concentrations of ePHF-tau over different periods of days in vitro (DIV) and for different durations (**Fig.1 a**). Our previous work mentioned that these rat neuronal cultures develop extensive interconnected 2D neuronal networks (**Fig. 1 b**). During this phase, neurite maturation takes place over the first three weeks, while excitatory synaptic maturation is established from the third week [20–22]. We found that prolonged exposure (14 days) to ePHF-tau on early maturing neurons (DIV7) resulted in a significant reduction in both cell density (**Fig. 1c**) and the extent of the neuritic network (**Fig. 1d**), even at a dose of 0.5uM. However, short-term treatment of mature neurons (DIV22), even at high doses of 1.5uM, did not appear to affect cell density (**Fig. 1e**) or neural network complexity (**Fig. 1f**). Subsequently, in order to assess the impact of PHF-tau on synaptic integration, we measured the density of presynaptic (puncta vGlut1+), postsynaptic (puncta PSD-95+) proteins, as well as excitatory active zones defined by the number of pairs of puncta vGlut1+ and PSD-95+ (**Fig. 1g**). Three days of ePHF-tau in the culture medium of mature neurons (DIV22) did not alter the densities of vGlut1+, PSD-95+ or the number of excitatory active zones (**Fig. 1g-j**). However, when cultures were subjected to the same treatment but at a later time (DIV25), we observed that PHF-tau induced a notable increase in the number of vGlut1+ puncta and PSD-95+ puncta, increasing the number of potentially active excitatory synapses (**Fig. 1g, k-m**).

To deepen our analysis of the impact of ePHF-tau on brain cells, We opted to complement our results with mixed neuron-glial 3D cultures (**Fig. 2a**). These mixed three-dimensional systems offer an organization closer to brain physiology, favoring faster electrical maturation and increased resilience to nerve agents, compared with pure 2D neuronal cultures (**Fig. 2b**) [20]. We observed that culture constitutes an assembly of neurospheroids enriched in neurons (NeuN) and astrocytes (GFAP), and endowed with neurites (NFL) interconnecting spheroids (**Fig. 2b**). ePHF treatment at a concentration of 0.5 μM does not affect the formation of neurospheroid (**Fig. 2c**), the density of neurons (**Fig. 2d**) and the amount of neurofibrillary fibers that compose them (**Fig. 2e**). Using confocal microscopy imaging and three-dimensional reconstruction of the neurospheroids, we quantified the volumes of vGlut1 and PSD-95 in the proximal neurites to the neurospheroids (**Fig. 2f-g**). We revealed that exposure to ePHF-tau for three days did not change PSD-95 density (**Fig. 2g-h**) or vGlut1 (**Fig. 2g,i**) volumes. However, the volume of overlap between vGlut1 and PSD-95 is significantly increased by ePHF treatment, suggesting an increase in the number of excitatory active synaptic sites (**Fig. 2g-j**). These results illustrate how exposure to ePHF-tau affects neurons differently depending on their stage of maturation and organiszation, reducing neurite density and complexity in immature neurons while increasing of excitatory active zone and potential excitotoxicity.

### Extracellular PHF-tau impacts the mitochondrial composition of synaptosomes

To better understand how ePHF-tau treatment could induce potential excitotoxicity, we isolated synaptosomes from 3D cultures 3 days after treatment with ePHF-tau and analyzed using proteomics. (**Fig. 3a**). Enrichment analysis revealed that most proteins were associated with synaptic terms, confirming the quality of the synaptosome isolation (**Fig. 3b**). ePHF-tau treatment affected numerous G.O. terms that were notably related to mitochondrial respiratory chain and regulation of mitochondrial membrane potential (**Fig. 3b**). Interestingly, consistent with our microscopy data (**Fig. 1-2**), another most differentially enriched G.O. terms were associated with intracellular transport and glutamate metabolism (**Fig. 3b**). In synaptosomes, we have observed that the amount of tau protein is highly correlated with the presence of mitochondrial protein from the inner membrane (I.M.) and intermembrane space (IMS), (**Fig. 3c**). Among these mitochondrial proteins found in synaptosomes we found a significant (8-fold) increase in the number of “central dogma” pathway proteins in the 19 IMS-associated proteins and a 4-fold increase in the number of proteins involved in “signaling” in the 40 mitochondrial matrix proteins (**Fig. 3e**). Together, these results suggest that treatment with ePHF-tau alone can induce significant activation of mitochondrial functions in synapses, which may be responsible for the increase in active zone and potential excitotoxicity.

### Mitochondrial monitoring revealed reversed and delayed effects of ePHF-tau on neurons and astrocytes

The primary challenge with the findings related to mitochondrial disruption at synapses when toxic substances are present extracellularly is the complexity of discerning whether this disruption is a direct effect on neurons, astrocytes, or a secondary response to the disturbance of either population. To improve this understanding, a continuous monitoring of the dynamic effects of ePHF-tau on mitochondrial function in different cell subpopulation may provide many clues. In this context, we employed lentivirus-mediated expression of MitoTimer and MitoGo-ATeam2 biosensors, which specifically target neurons or astrocytes [8,9]. We monitored the mitochondrial dynamics within neurites (**Fig. 4**) and astrocytic fine processes (**Fig. 5**) over a 72-hour posttreatment period using advanced fluorescence microscopy equipped for high-throughput acquisition and analysis. Observations were conducted at 2-hour intervals, amounting to approximately 35 timepoints. As previously described, each measured parameter is normalized with reference to its base state obtained in the first acquisition [9,21].

In neurites (Fig. 4), we revealed that treatment with ePHF-tau induced significant, rapid (from 10 h) and lasting (over 70 h) changes in neurite mitochondria (**Fig. 4a-k**). We observed a significant increase in mitochondrial ATP production (**Fig. 4d**) and mitochondrial oxidation (**Fig. 4f-g**). The primary contributor to mitochondrial oxidation (**Fig. 4e**) was an increase in the level of oxidized complexes (red color from Mitotimer) (**Fig. 4f**) rather than a reduction in mitochondrial biogenesis (green color from Mitotimer) (**Fig. 4g**). Moreover, ePHF-tau induced mitochondrial enlargement with a swollen morphology (**Fig. 4h-k**). Mitochondrial swelling in neurites is characterized by increased diameter, surface area, and length (**Fig. 4h-j**). Additionally, we observed a significant decrease in the shape factor of mitochondria (**Fig. 4k**), suggesting structural modifications with decreased convexity. Despite being administered in a single dose, the treatment progressively escalated toward increased stress conditions over time. In astrocytes processes (**Fig 5a-c**), we found no alterations in the mitochondrial redox state (**Fig 5d, e-g**) or mitochondrial morphology (**Fig 5d, h-k)** within the first 36 hours of treatment. However, significant deviations were observed beyond this period: mitochondrial processes display stronger fluorescence in the greens (**Fig. 5c, d**), corresponding to a decrease in the mitotimer ratio and suggesting increased mitochondrial turnover caused by increased biogenesis. Morphologically (**Fig. 5c, g**), despite the lack of significance, we observed a trend toward mitochondrial filamentation (fusion>fission). These results suggest that neurons are very rapidly sensitive to the effects of ePHF-tau, which will induce strong activation of the mitochondrial system and ultimately its functional disruption, whereas the effect on astrocytes is cocomitant with the neuronal problems and appears more as an increase in biogenesis and dynamics.

## Discussion

This study highlights the complexity of the toxic effect of extracellular tau aggregates on mitochondrial and synaptic function. This toxicity affects the mitochondria of neurons both rapidly and progressively, while the mitochondrial system of astrocytes does not appear to be immediately sensitive to these aggregated forms of tau but is stimulated a few days later. At the synaptic level, the effect of ePHF-tau on different synaptic behaviors induces an increase in neuronal excitability, potentially explained by cytotoxicity. Our study used Tau aggregates extracted from the brains of patients diagnosed with A.D. but standardized with tau 2N4R in monomeric form. Reverse and delayed astrocytic mitochondria responses to ePHF-tau treatment highlight cell-specific vulnerabilities and adaptation mechanisms to mitochondrial stress. Indeed, neurons show very rapid mitochondrial modification, which will lead to severe alteration in the long term. Astrocytes, on the other hand, show a significantly delayed response, synchronized with neuronal alterations, suggesting that the cells are rather reacting than suffering. These results underline the importance of considering the distinct roles and responses of different CNS cell types when exploring the pathophysiology of extracellular forms of tau.

Recent in-depth studies on the consequences of insoluble tau protein deposits in neurons, a characteristic marker of tauopathies, have highlighted the ability of these deposits to generate complex dysfunctions within various signaling pathways. These dysfunctions frequently translate into significant alterations in synaptic function, highlighting the major impact of these aggregates on fundamental neuronal mechanisms [22]. Indeed, soluble tau aggregates have been shown to significantly inhibit long-term depression (LTD) in the dorsal hippocampus of anesthetized rats, revealing their direct influence on synaptic plasticity [23]. The mechanism involves the binding of tau to synaptic vesicles via its N-terminal domain, which disrupts presynaptic functions, including synaptic vesicle mobility and release rate, effectively reducing neurotransmission [24]. Our results revealed that even nonhyperphosphorylated ePHF-tau can significantly increase synaptic active zones. In addition, our proteomic analyses revealed a notable reduction in proteins associated with the conversion of glutamate to GABA in synaptosomes, suggesting a decrease in inhibitory potential, a finding that could partly explain the observed imbalance in inhibitory potential. Previous work has also demonstrated that tau aggregates can alter the function of postsynaptic receptors, such as NMDA and AMPA receptors, contributing to the dysregulation of intracellular calcium homeostasis and increased neurotoxicity [25]. Furthermore, neuronal hyperexcitability and changes in presynaptic markers highlight a loss of balance between excitation and inhibition in neuronal networks [26], an imbalance potentially responsible for the cognitive and behavioral deficits observed in tauopathies [27]. Although the exact mechanisms underlying this synaptic disruption were not directly addressed in our study, our findings reinforce the idea that the effect of tau aggregates does not depend exclusively on the presence of hyperphosphorylated forms, and the strong association with mitochondrial proteins suggests that mitochondrial dysfunction could be one of the earliest signs of synaptic disruption.

The accumulation of tau protein in synapses plays a significant role in the disruption of mitochondrial function, contributing to the neuronal dysfunction characteristic of neurodegenerative pathologies such as A.D. Our study revealed that the mere presence of extracellular tau can significantly affect mitochondrial homeostasis in a very short time. Given the considerable size of ePHFs, our observations suggest two plausible hypotheses: first, specific fragments of tau, such as the N-terminal region, can detach and infiltrate synapses, rapidly influencing mitochondria; second, the aggregate itself can activate extracellular receptors that modulate mitochondrial homeostasis. Previous studies have shown that the truncated NH2 fragment of tau, 20-22 kDa in size, is mainly present in the mitochondria of cryopreserved synaptosomes from brains affected by A.D. This same Tau NH2 fragment has also been identified in other non-AD tauopathies, indicating its potential role in exacerbating synaptic degeneration in these diseases [28]. Our synaptosome analyses revealed a significant correlation between the amount of tau and the disruption of essential mitochondrial functions. However, our data show no specific enrichment of N-ter or C-ter fragments of human tau in synaptosomes following treatment, suggesting that the disruption is not solely due to these fragments. Research in mouse models has shown that mitochondrial distribution is gradually altered in neurites containing tau aggregates. It has also been established that aggregated tau can inhibit mitochondrial calcium efflux via the mitochondrial Na+/Ca2+ exchanger (NCLX), thus affecting neurons [29]. Our study highlights that the mere presence of ePHF is sufficient to induce a rapid and marked increase in ATP production by mitochondria in neurites, subsequently leading to extensive fragmentation and oxidation of mitochondria. These findings suggest potential underlying mechanisms by which neurons release Tau aggregates, raising crucial questions about the relevance of specific treatments currently in phase 2 and 3 clinical trials.

Astrocytes play an essential role in the evolution of tauopathies, influencing the progression of these diseases through their accumulation of tau and their ability to capture and diffuse pathological tau. Changes in their phenotype in the context of tauopathies can lead to dysfunctions that compromise their support of neurons or cause neurotoxicity [30]. Furthermore, specific expression of Tau 4R in mice expressing human tau has been shown to worsen seizure severity and increase phosphorylated Tau deposition without inducing neuronal or synaptic loss [31]. Our research revealed that only the accumulation of Tau 3R, not Tau 4R, in dentate gyrus astrocytes led to mitochondrial disruption, neuronal dysfunction, and memory deficits in mice[8]. Our recent studies also indicated that mitochondria from iPSC-derived astrocytes accelerate turnover and increase filamentation when exposed to extracellular vesicles from neurons that accumulate tau 1N4Rs, whereas they fragment and oxidize in response to those from neurons that accumulate tau 1N3Rs [9].

In our experiment with 2N4R tau aggregates on astrocytes, we also observed a significant increase in mitochondrial turnover and filamentation, although these changes took longer to appear as with 4R neuron-derived E.V.s. Initially, the mitochondrial system of astrocytes shows few changes in the first two days after exposure to PHF-tau 2N4R, suggesting that filamentation and increased mitochondrial turnover might indicate a proinflammatory profile [32,33]. This observation raises the hypothesis that astrocytes and their mitochondrial system might be less sensitive to these forms of 2N4R aggregates than to those present in or on extracellular vesicles. In this study, we did not explore the mechanisms underlying these effects, particularly the potential for capture by the LRP1 protein which is present in astrocytes and plays a crucial role in eliminating certain forms of amyloid aggregates. It is difficult to determine whether this cellular adaptation results from a response to distress in neighboring neurons or whether astrocytes have captured forms of tau that have activated them. Another hypothesis beside usual signal of stress transmission to glia, could be that neurons transfer dysfunctional mitochondria to astrocytes, as recently demonstrated in different brain injury contexts [34] and to microglia [35].

## Conclusion

This study highlights the variability of the response of the mitochondrial system to tau aggregates, emphasizing that this reaction depends on the cell type involved. This discovery could shed light on some of the obstacles encountered by clinical trials in their quest for drugs and antibodies capable of reducing the pathological burden of tau. This study also highlights the crucial importance of considering tau’s different forms and isoforms in the design of future preclinical drug trials. This perspective emphasizes the need for a more nuanced and personalized approach in the search for effective treatments for tauopathy diseases, reflecting the inherent complexity of these pathologies and the diversity of cellular responses they engender. Ultimately, our results reinforce the idea that the fight against tauopathies requires an in-depth understanding of the molecular mechanisms specific to each cellular context, paving the way for more targeted and potentially more effective therapeutic strategies.

## List of abbreviations

A.D.: Alzheimer’s disease
ePHF-tau: extracellular paired helical filaments of tau
FTD: Frontotemporal dementia
LV: Lentiviral Vector
NFT: Neurofibrillary tangles of tau
PHF-tau: paired helical filaments of tau.
PSP: progressive supranuclear palsy
ROI: Region of interest
VOI: Volume of interest

## Acknowledgments

The authors thank Maria Rey and Nicole Déglon, Lausanne University Hospital (CHUV) and University of Lausanne (UNIL), Department of Clinical Neurosciences (DNC), and Laboratory of Cellular and Molecular Neurotherapies (LCMN) for their help in the production of lentiviral vectors.

## Author contributions

Study design and conceptualization: K.R. Writing, review, and editing: K.R, V.Z. Support for manuscript editing: N.P. Experimentation: E.P, V.Z, A.B, Y.V, K.F. Funding acquisition: K.R, F.C. Supervision: K.R. All authors have read and agreed to the published version of the manuscript.

## Ethics declarations

For murine-derived primary cultures, experiments were performed according to an ethics protocol approved by our institutional review committee CER-VD 2018-01622, Lausanne, Switzerland.

